# Sex-segregated range use by black-and-white ruffed lemurs (*Varecia variegata*) in Ranomafana National Park, Madagascar

**DOI:** 10.1101/2020.03.30.016774

**Authors:** Andrea L. Baden, Brian D. Gerber

## Abstract

Ranging behavior is one important strategy by which nonhuman primates obtain access to resources critical to their biological maintenance and reproductive success. As most primates live in permanent social groups, their members must balance the benefits of group living with the costs of intragroup competition for resources. However, some taxa live in more spatiotemporally flexible social groups, whose members modify patterns of association and range use as a method to mitigate these costs. Here, we describe the range use of black-and-white ruffed lemurs (*Varecia variegata*) at Mangevo, an undisturbed primary rainforest site in Ranomafana National Park, Madagascar and characterize sex-differences in annual home range area, overlap, and daily distances traveled. Moreover, we characterized seasonal variability in range use and ask whether ranging behaviors can be explained by either climatic or reproductive seasonality. We found that females used significantly larger home ranges than males, though sexes shared equal and moderate levels of home range overlap. Overall, range use did not vary across seasons; though within sexes, male range use varied significantly with climatic variation. Moreover, daily path length was best predicted by day length, female reproductive state, and sex. While the patterns of range use and spatial association presented here share some similarities with ‘bisexually bonded’ community models described for chimpanzees, we argue that ruffed lemurs represent yet another community model wherein ‘neighborhoods’ of overlapping males and females affiliate with each other moreso than with other, more distantly located community members, and to the exclusion of members from adjacent communities.

## INTRODUCTION

In many primates, individuals live, travel, and forage together in cohesive units or “social groups.” Despite exhibiting variable preferences for social partners [Cords, 2002; Silk et al., 2006b; Silk et al., 2006a; Schülke et al., 2010; Seyfarth et al., 2014; Perry, 2013], resource utilization [Boinski, 1988; Fragaszy, and Boinski, 1995; King et al., 2009], and/or spatial position within the group [Janson, 1990a; Janson, 1990b; Ron et al., 1996; Hall, and Fedigan, 1997]; reviewed in [Hirsch, 2007], patterns of individual movement in these taxa are generally broadly coordinated, and range use — including home range size, overlap, and daily path length (DPL) — is more or less coincident amongst group members (e.g., [Strandburg-Peshkin et al., 2015]. As such, there is expected to be little variation in individual ranging patterns among age-sex classes within cohesive social groups (though these patterns break down around natal dispersal and secondary transfer events (e.g., [Jack, and Isbell, 2009] and references therein)). By contrast, in some taxa, group members are able to individually optimize the costs and benefits of group living, and groups are much less spatiotemporally constrained. In these taxa, members of socially and geographically circumscribed groups (or “communities”) associate in temporary, flexible subunits (i.e., “parties” or “subgroups”) that vary in size, cohesion, membership, and duration via a strategy known as “fission–fusion” (*sensu* [Kummer, 1971]; reviewed in [Aureli et al., 2008]. Such behavioral flexibility allows group members to individually adjust their patterns of association in accordance with ecological and social constraints [Williams et al., 2002; Symington, 1990; Chapman et al., 1995; Lehmann, and Boesch, 2004][Mitani et al., 2002] and references therein), variation which is in turn reflected in their corresponding patterns of range use (reviewed in [He et al., 2019].

In chimpanzees and spider monkeys—the best-studied of the non-human primates exhibiting high fission-fusion dynamics—patterns of social association and range use are generally sex-segregated (e.g., [Goodall, 1986; Symington, 1990; Hasegawa, 1990; Chapman, and Wrangham, 1993; Williams et al., 2002]; but see [Lehmann, and Boesch, 2005; Lehmann, and Boesch, 2008] for discussion of bisexually-bonded patterns of chimpanzee fission-fusion dynamics). Males tend to be more gregarious than females (e.g., [Otali, and Gilchrist, 2006; Lehmann, and Boesch, 2008]; [Wrangham, 2000]), and exhibit stronger social bonds with each other than either mixed-sex or female-female dyads [Machanda et al., 2013; Fedigan, and Baxter, 1984; Gilby, and Wrangham, 2008; Shimooka, 2003; Slater et al., 2009]; but see [Lehmann, and Boesch, 2008; Williams et al., 2002]. By contrast, females—particularly those with infants or dependent offspring— are more often found alone or in small same-sex parties [Nishida, 1968; Wrangham, and Smuts, 1980; Goodall, 1986; Symington, 1988a; Symington, 1990; Chapman, 1990; Lehmann, and Boesch, 2008; Wrangham, 2000].

Because individual movement in these taxa is generally less constrained, sex biases in social association are typically reflected in sex-segregated patterns of range use [Wrangham, and Smuts, 1980; Stumpf, 2007]. Males tend to travel longer daily distances [Wrangham, and Smuts, 1980; Doran, 1997; Wallace, 2008]; but see [Herbinger et al., 2001], use larger overall home ranges [Wrangham, and Smuts, 1980; Symington, 1988b; Symington, 1990; Chapman, and Wrangham, 1993; Williams et al., 2002; Shimooka, 2005; Wrangham et al., 1992; Nunes, 1995], and spend more time in the peripheries of their territory than do females [Chapman, and Wrangham, 1993; Chapman, 1990; Lehmann, and Boesch, 2005; Mitani, and Watts, 2005; Shimooka, 2005; Wallace, 2008]. Males also tend to share larger, more overlapping home ranges with other males and most, if not all females within their community [Chapman, and Wrangham, 1993; Shimooka, 2005; Symington, 1988b; Nunes, 1995; Wrangham, 1979]. Females in these same communities do not typically utilize their entire range, and instead restrict their movement to smaller, more central areas within the territory [Wrangham, and Smuts, 1980; Chapman, and Wrangham, 1993; Williams et al., 2002; Shimooka, 2005]; but see [Chapman, 1990; Boesch, and Boesch-Achermann, 2000; Wallace, 2008; Lehmann, and Boesch, 2005]. Together, these patterns have been characterized by some as following a ‘male-bonded’ community model [Lehmann & Boesch 2005; Fig. 1].

**Figure 1.**
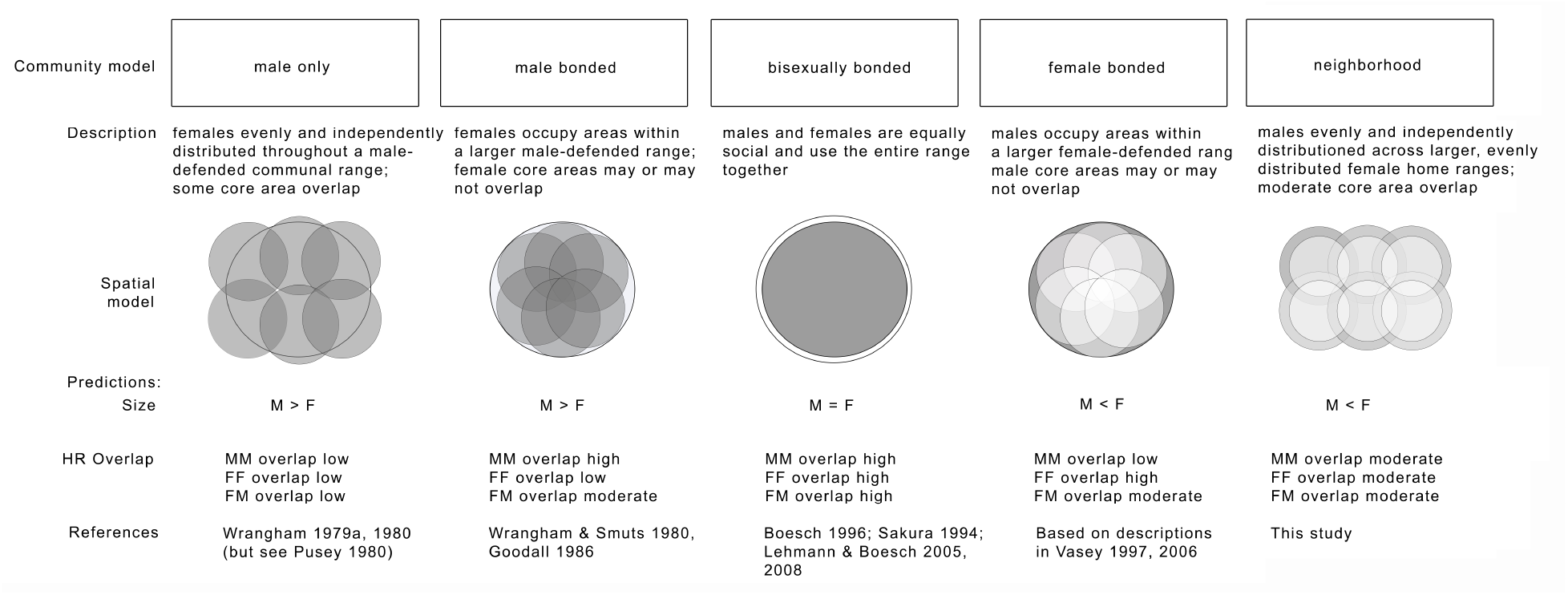
Though traditionally used to characterize chimpanzees, community models of fission-fusion dynamics allow for clear predictions regarding home range (HR) size, overlap, and the spatial distribution of males and females in any nonhumanprimate.

There is, however, notable inter- and intra-population variation in the patterns of association and range use observed in these species. For instance, some populations of chimpanzees (e.g., Bossou and Taï in West Africa) and spider monkeys (e.g., Yasuni) can be characterized as exhibiting a ‘bisexually-bonded’ community model, wherein males and females are equally gregarious, spending more time in mixed-sex parties [Boesch, 1991], and exhibiting more similar range use and overlap [Sugiyama, 1988; Sakura, 1994; Boesch, 1996, Lehmann and Boesch, 2005; Lehmann and Boesch, 2008; Spehar et al., 2010](Fig. 1), patterns that have been attributed to higher resource availability, lower population size/density, and reduced predation and/or anthropogenic pressure relative to eastern sites (reviewed in [Lehmann, and Boesch, 2005]. Moreover, intra-population variation has been attributed to inter- and intra-annual seasonality; in fact, many argue that fission-fusion dynamics may have primarily evolved in response to shifting resource availability, allowing individuals to maximize the benefits of group living (e.g., predator avoidance, resource defense) while minimizing within-group resource competition [Boesch, and Boesch-Achermann, 2000](but see [Lehmann et al., 2007]. During periods of resource scarcity, individuals typically form smaller, more dispersed foraging parties (e.g., [Itoh, and Nishida, 2007; Aguilar-Melo et al., 2018; Aureli et al., 2008; Anderson et al., 2002; Boesch, 1996; Chapman et al., 1995; Doran, 1997; Mitani et al., 2002; Shimooka, 2003; Symington, 1990; Symington, 1988a; Newton-fisher et al., 2000; Matsumoto-Oda et al., 1998] Ghiglieri, 1984; Klein & Klein, 1977; Wrangham, 1977, 1980; Wrangham et al., 1992, 1996; but see [Boesch, 1996; Hashimoto et al., 2003; Hashimoto et al., 2001; Lehmann, and Boesch, 2004; Riedel et al., 2011]; Stanford et al., 1994). And while there is mixed evidence of seasonal changes in overall territory size (e.g., [Basabose, 2005; Nakamura et al., 2013; Moore et al., 2018], animals will commonly adjust habitat use seasonally by traveling farther [Moore et al., 2018]-Mayeve; but see [Doran, 1997; Herbinger et al., 2001; Moore et al., 2018]-Cyamudongo) or into rarely used parts of their range (e.g., [Basabose, 2005; Goodall, 1986; Herbinger et al., 2001].

While the vast majority of primate fission-fusion literature has traditionally focused on chimpanzees and spider monkeys, there is growing evidence that fission-fusion dynamics are, in fact, more common than previously recognized (e.g., [Aureli et al., 2008]), and that the demographic, ecological, and social variables described above can be applied more broadly to explain fission-fusion dynamics across the primate order. For instance, in many nonhuman primates, group fissions and concomitant changes in ranging behavior commonly occur within a feeding context (e.g., [Dolado et al., 2016; Dolado et al., 2017; Izar et al., 2012; Ren et al., 2012] and/or in accordance with social variables such as subgroup size or the presence of mates (e.g., [Ellis, and Di Fiore, 2019; Dias, and Strier, 2003; Strier, 2018]. Here, we add to this growing body of literature by quantifying patterns of range use and overlap in a strepsirrhine with high fission-fusion dynamics, the black-and-white ruffed lemur (*Varecia variegata*) [Baden et al., 2016].

Ruffed lemurs are relatively large-bodied (2.5– 4.8 kg; [Baden et al., 2008]), arboreal frugivores endemic to the eastern rainforests of Madagascar [Baden, 2011; Balko, 1998; Morland, 1991a; Rigamonti, 1993; Ratsimbazafy, 2002; Vasey, 2000]. Ruffed lemurs live in large social groups (e.g., 11 to 31 individuals) that are characterized by high levels of fission–fusion social dynamics [Holmes et al., 2016b; Baden, 2011; Morland, 1991a; Rigamonti, 1993; Baden et al., 2016; Vasey, 1997; Morland, 1991b]; but see [Balko, 1998; Ratsimbazafy, 2002]. Aspects of their social behavior, including subgroup size, composition, and cohesion vary by season, resource availability, and female reproductive state [Morland, 1991a; Rigamonti, 1993; Vasey, 2006; Baden et al., 2016; Morland, 1991b]. While there has been a recent uptick in studies investigating ruffed lemur fission-fusion dynamics (e.g., [Holmes et al., 2016a; Baden et al., 2016; Baden et al. in press], those characterizing ruffed lemur range use within this same context remain limited and provide incongruous results. For instance, one study of red ruffed lemurs (*Varecia rubra*) from the Masoala Peninsula described females as utilizing large, seasonally variable home ranges that overlapped extensively with other females, and which encompassed the smaller, non-overlapping territories of males [Vasey, 2006]. Despite disparate patterns of home range area between sexes, daily distances traveled did not vary between sexes, except within the context of mating, when males traveled significantly farther than females. By contrast, Morland’s [Morland, 1991b] study of *V. variegata* on Nosy Mangabe described males as ranging farther (i.e., using larger annual home ranges that overlapped with multiple adjacent communities) and traveling significantly longer average daily distances than females. While these early results stand in stark contrast, previous research has also noted substantial inter-population variability in other aspects of their demography and social behavior, including population density, social organization, and affiliative interactions (reviewed in [Baden et al., 2016], suggesting that range use may be equally as variable.

Because shared range use provides individuals the opportunity for repeated social interactions (e.g., [Clutton-Brock, 1989; Kossinets, and Watts, 2006], it is critical that studies consider individual patterns of range use and overlap when investigating the evolution of social behavior, particularly within taxa exhibiting high fission-fusion dynamics (e.g.,[Best et al., 2014; Carter et al., 2013; Frère et al., 2010; Strickland et al., 2014; Baden et al.]. Thus, to broaden our understanding of ruffed lemur social behavior, including patterns of social association and fission-fusion dynamics, we must also improve our understanding of ruffed lemur range use. To this end, we characterize annual patterns of range use, including home range area, overlap, and daily path length within one *V. variegata* community. Further, we evaluate whether and how ruffed lemur range use varies with seasonal shifts in ecology and reproductive physiology.

Based on earlier findings in this and other non-human primate species, we hypothesize that (H1) patterns of ruffed lemur range use will vary in accordance with climatic season (and thereby resource availability), as do their seasonal patterns of fission-fusion social dynamics (*sensu* Baden et al. 2016). Because resource competition is presumed to be equitable among males and females in this species, we predict that (P1.1) sexes will not differ in their range use in across climatic seasons. We predict that both males and females will (P1.2) travel farther, and have (P1.3) larger, (P1.4) more overlapping ranges during periods of high resource availability (e.g., warm-wet seasons). By contrast, in accordance with the time minimizing strategies of many lemur species (*sensu* [Wright, 1999; Beeby, and Baden], we further predict that ruffed lemurs will (P1.5) travel less (i.e., shorter daily travel distances), and adopt home ranges that are (P1.6) smaller, and (P1.7) more exclusive (i.e., less spatial overlap) during cool, wet periods when resources are scarce. Finally, because cool-dry periods align with rising resource abundance, we predict that (P1.8) home range size, (P1.9) overlap, and (P1.10) daily distances traveled will be intermediate to those exhibited during either warm-wet or cool-dry periods.

Alternatively, because patterns of resource availability are largely, though imprecisely concordant with patterns of ruffed lemur reproduction, we hypothesize that (H2) range use may instead reflect the unique mating strategies of the species [Baden, 2011; Baden et al., 2016]. In this regard, we predict that (P2.1) males and females will differ in their patterns of range use dependent on female reproductive state. We predict that males will (P2.2) travel more, and use ranges that are (P2.3) relatively larger and (P2.4) more overlapping with females during the brief mating and subsequent gestation season than during either nonbreeding or lactation seasons (i.e., mating/gestation > nonbreeding or lactation). By contrast, we predict that females will (P2.5) travel relatively longer distances and utilize (P2.6) larger (P2.7) more overlapping ranges when unconstrained by infants (i.e., nonbreeding > gestation or lactation). We further predict that (P2.8) female home range size and (P2.9) daily distance traveled will be most constrained (i.e., smallest) during the period of lactation and high infant dependence, but that their (P2.10) home ranges will overlap more with other females during lactation than mating/gestation due to their communal créching infant care strategy [Baden et al., 2013a; Baden, 2019].

## METHODS

### Study site & subjects

We collected data from one black-and-white ruffed lemur (*Varecia variegata*) community at Mangevo bushcamp in Ranomafana National Park, Madagascar (RNP) during 16 months (August - December 2007; February - December 2008). Mangevo (21°22’60”S, 47°28’0”E) is a mid-elevation (660 - 1,200m) primary rainforest site within the southeastern parcel of RNP, 435 km^2^ of continuous montane rainforest located in the southeastern escarpment of Madagascar’s central high plateau (21°02’– 21°25’S and 47°18’– 47° 37’E; [Wright et al., 2012]).

At the time of this study, the community included 24 adults and subadults (8 adult females, 11 adult males, 5 subadult males). Nineteen infants were born in the 2008 birth season and were present from October - December 2008. Of the adults, 5 females and 3 males were radio-collared and targeted for regular follows. Individuals with collar-tags (but no radio-collar, n = 16) were opportunistically targeted for focal follows. Sampling efforts resulted in a total of 4,044 focal observation hours.

## Data Collection

### Behavioral monitoring

Two teams of four observers each conducted dawn-to-dusk follows on focal individuals, (i.e., two animals were followed daily). A focal animal was located at the beginning of each observation period via radio-telemetry. Only independent individuals (adults and subadults) were targeted for follows. New focals were selected daily. Focals were never sampled on consecutive days and every effort was made to follow all subjects at least once per month. If an individual with a collar-tag was located in association with a radio-collared focal individual prior to 10:00 h, this individual became the new focal for that observation period. Observation periods ranged in duration between 8 to 11 hours depending on seasonal differences in day length and time needed to locate animals at dawn.

Upon initial contact with the focal individual, we recorded the number and identities of all other individuals present within the subgroup. Subgroups were defined as all independent individuals (i.e., adults and subadults) within a 50 m radius of the approximate subgroup center that exhibited coordinated behavior and travel (see [Baden et al., 2016] for details). After initial contact, we monitored subsequent changes in subgroup size, composition (age/sex class, individual identity), and cohesion (i.e., the greatest distance between any two subgroup members), as well as activity state of the focal subject using instantaneous scan sampling techniques collected at 5-min intervals [Altmann, 1974].

We collected simultaneous data on subgroup location from the approximate group center at 10-minute intervals using a handheld Garmin^®^ HCx GPS unit. Spatial coordinates were recorded only if estimated positional error was less than 10 m.

### Ethical note

We used remote anaesthetization techniques to capture and collar study subjects following established protocol [Glander, 1993]. Animal captures were performed by a team of skilled Malagasy technicians (Madagascar Biodiversity Partnership) in the presence of at least one trained veterinarian per capture season (Randy Junge, Felicia Knightly, Edward Louis, Angie Simai). A Dan-Inject (Brrkop, Denmark) Model JM CO_2_-powered rifle was used to administer 10mg/kg estimated body weight of Telzaol^®^ (Fort Dodge, LA) via 9 mm (3/8”) Type ‘P’ disposable Pneu-darts^™^ (Williamsport, PA). GPS coodinates and trail headings were collected at the site of each capture, after which animals were transported back to camp for processing. During processing, subjects were monitored for heart rate, respiratory rate, and body temperature and were given a subcutaneous balanced electrolyte solution (LRS, lactate ringer solution). Subjects were then allowed to recover from anesthesia in breathable fabric bags (~3 h) and then released at the site of capture. Any subjects who had not recovered prior to dusk were kept overnight and released at the capture site the following morning. Upon release, subjects were followed to ensure full mobility in trees and to confirm that there were no injuries as a consequence of the captures. All collars were well below the 5% threshold of the subjects’ weight recommended for arboreal animals and were not observed to impede our subjects’ normal behaviors.

Research protocols were in compliance with and permission granted by Stony Brook University IACUC #2005-20081449 and Madagascar’s National Parks (ANGAP/MNP), and adhered to the American Society of Primatologists (ASP) Principles for the Ethical Treatment of Non-Human Primates.

### Data Analyses

We performed home range analyses with Home Range Tools [Rodgers et al., 2007] add-on for ArcGIS (ESRI, Redlands, CA). We did not subsample ranging data, as this has been shown to reduce the accuracy and precision of home range estimates ([De Solla et al., 1999; Blundell et al., 2001; Fortin, and Dale, 2005; Fieberg, 2007]. We did, however, limit our analyses to only those spatial coordinates for which corresponding data on subgroup size and composition were available (n=21,748 points). From this dataset, we calculated the communal home range (i.e., the home range used by all members of the focal ruffed lemur community), as well as individual home ranges for each of its members. Individual home ranges were calculated using location points collected while an individual was the subject of focal observations, as well as when the individual was a member of the subgroup being followed (i.e., was recorded as present during a focal follow of another individual).

Certain individuals were difficult to locate and were peripheral to social interactions within the community. Because home range analyses should use data that encompass the full range of movement behavior exhibited by an animal [Harris et al., 1990], data from infrequently encountered animals may not be representative of their true movement behaviors. We therefore investigated the relationship between sample size and measures of both annual and seasonal home range area with an incremental area analysis (i.e., increment plots; [Kenward, 2001; Kenward, and Hodder, 1996], whereby we iteratively added spatial locations and estimated home range size for each individual to evaluate asymptotic stability in our estimates. This was done for the communal territory, as well as for individual territories within the larger communal range. We found that communal territory stabilized after ~500 fixes. Annual individual home range estimates tended to asymptote earlier, after approximately 250 location points. To account for monthly variation in sampling effort, we omitted animals with fewer than 250 location points spread across 25 sampling days throughout the year from annual home range analyses. It is important to note that, in a majority of cases, sampling effort was evenly distributed throughout the year. Thus, while 250 location points was a minimum criterion for infrequently encountered animals, most individuals included in the analysis were sampled at least twice as often. Similarly, estimates of seasonal home range area reached asymptotes around 100 location points, and thus to ensure representative sampling, individuals with fewer than 100 location points spread across 10 sampling days within a given season were omitted from further analyses.

### Annual range use and overlap

Using the datasets described above, we estimated range use in two ways: using kernel density estimates (KDEs) and minimum convex polygons (MCPs). The first method, kernel density estimates (KDE) are widely regarded as a robust probabilistic estimator for making inferences about home range size, as well as about patterns of use within the home range (utilization distribution: [Worton, 1989; Powell, 2000]). KDEs were calculated using a bivariate normal distribution, rescaling X-Y coordinates to unit variances as recommended by [Silverman, 1986]. Raster cell size was set to 10 x 10 m to reflect the spatial resolution of our data. Kernel home range estimators are sensitive to the choice of smoothing parameter [Silverman, 1986]; this is especially true with large datasets and when animals exhibit strong site fidelity [Hemson et al., 2005]. Given our large dataset and observations that animals regularly transverse the same areas, we used the root-n smoothing parameter, as it has been found to overcome these issues [Steury et al., 2010]. Home range size was estimated for the community as a whole, as well as for all individuals separately, and ranges were evaluated using 95% kernel isopleths.

The second method, minimum convex polygons (MCPs), creates individual polygons that include all locations where a particular individual was recorded [Mohr, 1947]. However, the method suffers from sample size effects and is greatly affected by outliers, such that MCP estimates often contain large areas never used by an animal [Laver, and Kelly, 2008; Powell, 2000]. MCP estimates might also affect small-scale comparisons, i.e., within species or populations, although when large differences occur most of the variation is due to real differences [Nilsen et al., 2008]. This method was used exclusively to facilitate comparison with previous studies of ruffed lemurs. To help mitigate outlier effects, we calculated 95% MCPs via the fixed mean, a method employed to control for rare but observed excursions outside of the home range.

In addition to home range area, we also generated estimates of home range overlap. Kernel overlap was calculated using a utilization distribution overlap index (UDOI; [Fieberg, and Kochanny, 2005]) implemented in the R package adehabitat [Calenge, 2006]. The UDOI is an index of space-use sharing between two utilization distributions. UDOI values can range from 0 to 1, with a UDOI of 0 indicating no home range overlap and a UDOI of 1 indicating that home ranges are uniformly distributed (i.e., 100% overlap). Values can also be >1 if both UDs are nonuniformly distributed and also have a high degree of overlap. Values <1 indicate less overlap relative to uniform space, whereas values >1 indicate higher than normal overlap relative to uniform space. We calculated annual and seasonal UDOIs for all pairs of individuals included in our study. MCP overlap in individual home ranges was calculated as the proportion of shared area between two polygons. MCP overlap was calculated for communal home range area and individual annual home range areas only.

### Seasonal range use

We consider a home range as “the space which the animal both uses and traverses” during an ecologically meaningful time period [Burt, 1943]. Thus, to further investigate patterns of ruffed lemur spatial ecology, we also calculated seasonal measures of individual home range area and overlap. Specifically, we used data on climate, phenology, and female reproductive state to separate months into discrete seasons with intervening transitional periods (see [Baden, 2011; Baden et al., 2016] for detailed methods), and then analyzed data in accordance with the *climatic* or *reproductive season* in which they were collected.

Measures of fruiting seasonality and climate were closely associated [Baden et al., 2016]. We therefore used natural breaks in the data to categorize ranging coordinates as falling into one of three *climatic seasons* (Warm-Wet/Peak Fruiting: November 2007 – February 2008; Cool-Wet/Fruit Scarcity: May-July 2008; and Cool-Dry/Low-to-Moderate Fruiting: August-September 2007, 2008; March, April, and October were considered transitional seasons and were excluded from this analysis), as well as one of three *reproductive seasons* (Nonbreeding: August 2007-June 2008; Gestation: July-October 2008; Lactation: November-December 2008) (Table 1; see also Fig. 1 in [Baden et al., 2016]).

**Table 1.**
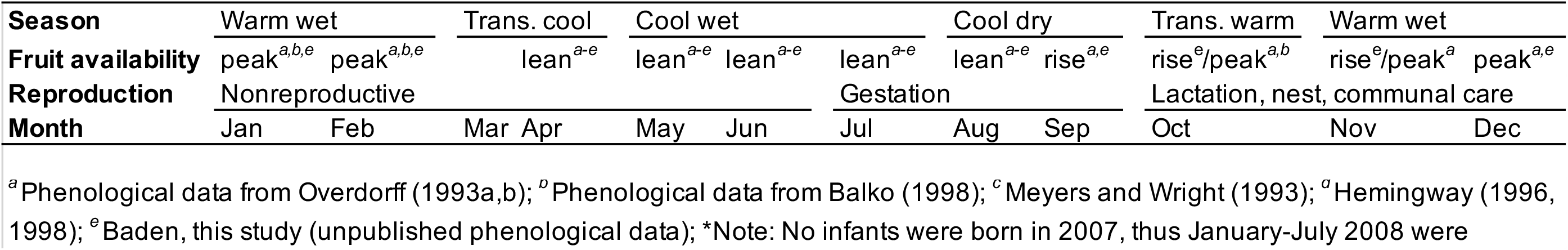
Patterns of climatic, phenological, and ruffed lemur reproductive seasonality in Ranomafana National Park, Madagascar.

### Daily range use

Finally, to gain a finer-scale understanding of daily range use, we calculated daily path lengths (DPL) for each individual as the sum of the Euclidean distances between consecutive GPS coordinates taken every 10-minutes. We included only full-day follows of focal individuals for which locations were recorded completely between morning and evening sleep trees, or data collection started prior to 0700 h with ≥ 9 subsequent hours of observation and fewer than 5% missing observations.

### Statistical analyses

All analyses were conducted in R version 3.6.2 (R Core Team 2019). Alpha was set *a priori* at α = 0.05. Individual home range area and overlap were statistically compared by sex or subgroup type and season using a combination of nonparametric Skillings-Mack and Mann Whiney-U tests. Daily path lengths were analyzed using a generalized linear mixed-effects model in the lme4 package [Bates et al., 2014] in R. Fixed effects included sex, day length, mean daily rainfall, a categorical classification of climatic season, a categorical classification of reproductive state, and the presence and number of infants. To explicitly account for individual variation in DPL, we included Individual ID as a random effect. Using additive and interactive effects of our variables of interest, we *a priori* constructed biologically driven models; we evaluated model parsimony using Akaike’s Information Criterion with a small sample size bias correction (AICc). To incorporate model selection uncertainty, we model-averaged all parameter estimates [Burnham, and Anderson, 2002].

## RESULTS

### Annual range use

Home range estimates for the entire Mangevo ruffed lemur community were between 87 ha (KDE) and 120 ha (MCP). Within the community, individual annual home ranges varied between 11.5 and 20.6 ha. Females (n=5) used significantly larger annual home ranges than did males (n=7) (MCP: Mann-Whitney *U* = 10.0, *P* = 0.04; Kernel: Mann-Whitney *U* = 9.0, *P* = 0.03; Table 2). On average, female home ranges were estimated to encompass 16.9 ha ± 1.74 SD (KDE) to 26.3 ha ± 4.50 SD (MCP), whereas an average male’s home range was estimated to cover 13.04 ha ± 0.98 SD (KDE) to 17.5 ha ± 1.22 SD (MCP)(Fig. 2).

**Figure 2.**
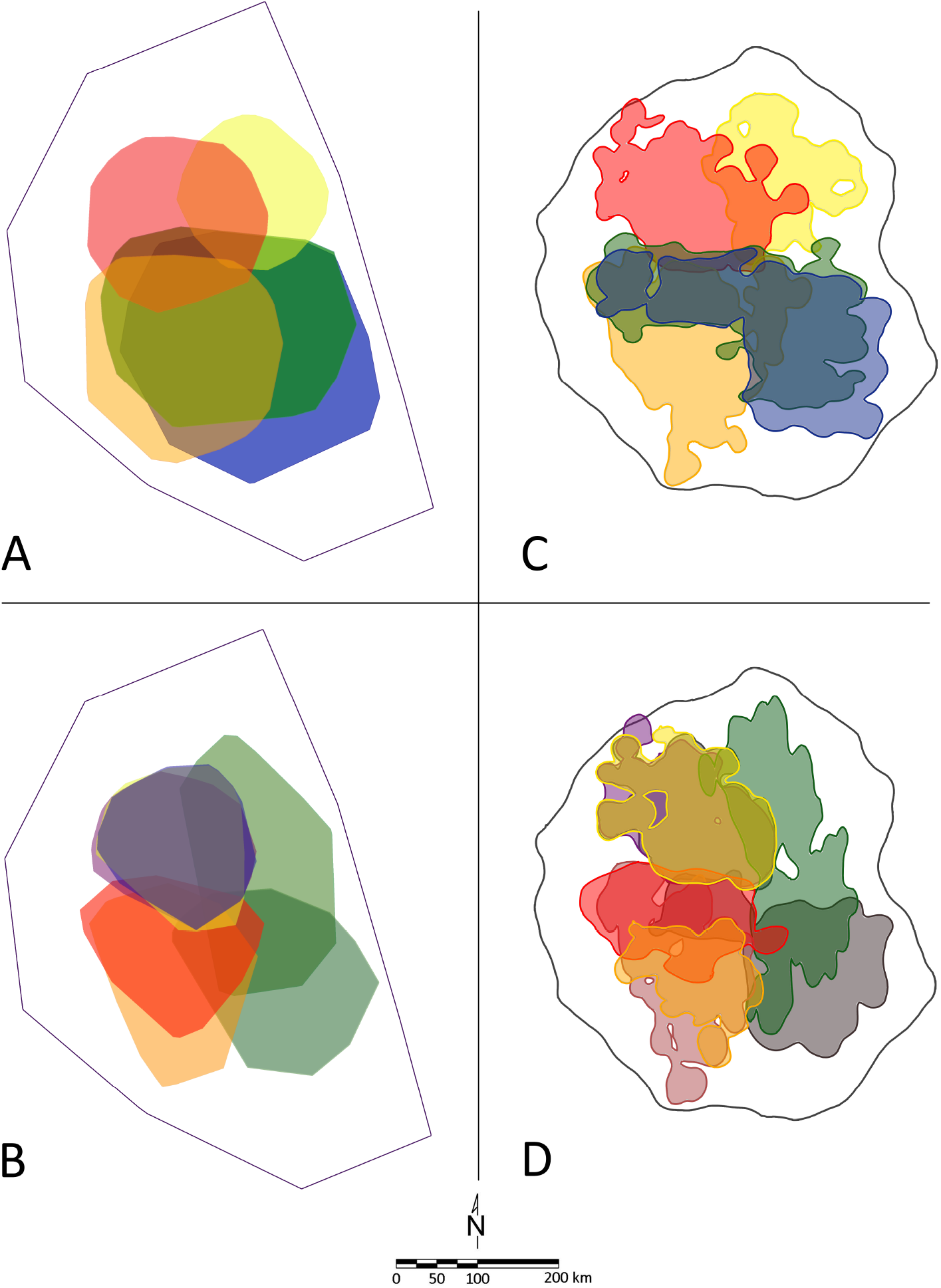
Differences in annual home range size and overlap for females (A & C, top, n=5) and males (B & D, bottom, n=8) using minimum convex polygon (MCP: A & B, left) and 95% kernel density estimates (KDE: C & D, right).

**Table 2.** Individual annual home range area and overlap. Females used significantly larger home ranges than males (MCP: Mann-Whitney *U* = 10.0, *P* = 0.04; Kernel: Mann-Whitney *U* = 9.0, *P* = 0.03). However, sexes did not differ in their degree of home range overlap. Analysis includes individuals with ≥ 25 annual sampling days.

| Collar ID | Sex | Located days | # location points | 95% MCP (ha) | 95% Kernel (ha) |
| --- | --- | --- | --- | --- | --- |
| RADIO-BLUE (rB) | F | 90 | 4277 | 38.7 | 20.6 |
| RADIO-BLUE-GREEN (rG) | F | 107 | 4578 | 32.9 | 19.4 |
| RADIO-ORANGE (rO) | F | 75 | 3386 | 26.6 | 18.7 |
| RADIO-RED (rR) | F | 92 | 3960 | 19.6 | 14.1 |
| RADIO-BLUE-YELLOW (rY) | F | 72 | 3640 | 13.6 | 11.5 |
| Female Mean |  | 87.2 | 3968.2 | 26.3 | 16.9 |
| BLACK-GREEN (BG) | M | 48 | 808 | 22.8 | 17.0 |
| NO COLLAR (NC) | M | 49 | 735 | 18.3 | 13.8 |
| RADIO-BLACK-GREEN (rBG) | M | 74 | 2163 | 20.6 | 16.1 |
| RADIO-PURPLE-SILVER (rPS) | M | 76 | 2403 | 14.8 | 12.3 |
| BLACK-BLUE (BB) | M | 58 | 1792 | 14.9 | 12.0 |
| RED-GREEN (RG) | M | 58 | 1276 | 15.8 | 12.4 |
| YELLOW-PURPLE (YP) | M | 56 | 1607 | 15.0 | 12.7 |
| Male Mean |  | 59.9 | 1540.6 | 17.5 | 13.8 |
| $p$ | | | | 0.04 | 0.03 |

| Sex-Sex | N dyads | Overlap |  |
| --- | --- | --- | --- |
|  |  | 95% MCP | 95% Kernel |
| Female-Female | 10 | 0.41 ± 0.09 | 0.16 ± 0.05 |
| Male-Male | 21 | 0.35 ± 0.07 | 0.24 ± 0.09 |
| Female-Male | 35 | 0.40 ± 0.05 | 0.29 ± 0.07 |
| $p$ | | 0.62 | 0.70 |

Neither sex used the entire communal home range. Rather, males and females concentrated their total annual space-use to only a proportion of the larger communal territory. Females used, on average, between 19.2% – 21.8% of the total community home range (KDE: mean % of communal range = 19.2%, n = 5, range = 13.1 – 23.5%; MCP: mean % of communal range = 21.8%, n = 5, range = 11.3 – 32.2%), while males used between 11.5% – 15.7% (KDE: mean % of communal range = 15.7%, n = 7, range = 13.7 – 19.4%; MCP: mean % of communal range = 11.5%, n = 7, range = 10.0 – 14.1%).

Across same-sex and mixed-sex dyads, home range overlap was low to moderate (KDE: 16 – 29%; MCP overlap: 35 – 41%) and sexes did not differ significantly in their proportion of home range overlap (Table 2; Fig. 2).

### Seasonal range use

Overall, individual home range size did not vary significantly across climatic seasons (Skillings-Mack test, SM = 4.87, df = 2, *P* = 0.09), although patterns of range use did differ between sexes (Table 3; Fig. 3). Within sexes, female home range size did not vary significantly by climatic seasons (Skillings-Mack test, SM = 3.60, df = 2, *P* = 0.17). By contrast, males exhibited marked seasonal variability in home range size (Skillings-Mack test, SM = 8.14, df = 2, *P* = 0.02). Males used significantly smaller home ranges during the cool-wet season than either the cool-dry season (Wilcoxon Signed Ranks, Z = 28.0, *P* = 0.02), or the warm-wet season (Wilcoxon Signed Ranks, Z = 15.0, *P* = 0.04)(Fig. 3). Male home ranges did not differ significantly between cool-dry and warm-wet seasons (Wilcoxon Signed Ranks, Z = 10.0, *P* = 0.50). Further, male and female home range size differed in two of three climatic seasons: males used significantly smaller home ranges than females during the cool-wet (females = 14.59 h, males = 6.42 ha; Mann-Whitney U = 3.0, *P* = 0.02) and warm-wet seasons (females = 16.38 ha, males = 9.64 ha, Mann-Whitney U = 3.0, *P* = 0.05; Table 3). Home range areas were similar in size only during the cool-dry season, when female range use decreased (females = 11.84 ha, males = 10.69 ha, Mann-Whitney U = 13.0, *P* = 0.53; Table 3). We found no significant difference in home range overlap across seasons among same- and mixed-sex dyads (Skillings-Mack test, SM = 2.05, df = 2, *P* = 0.36).

**Figure 3.**
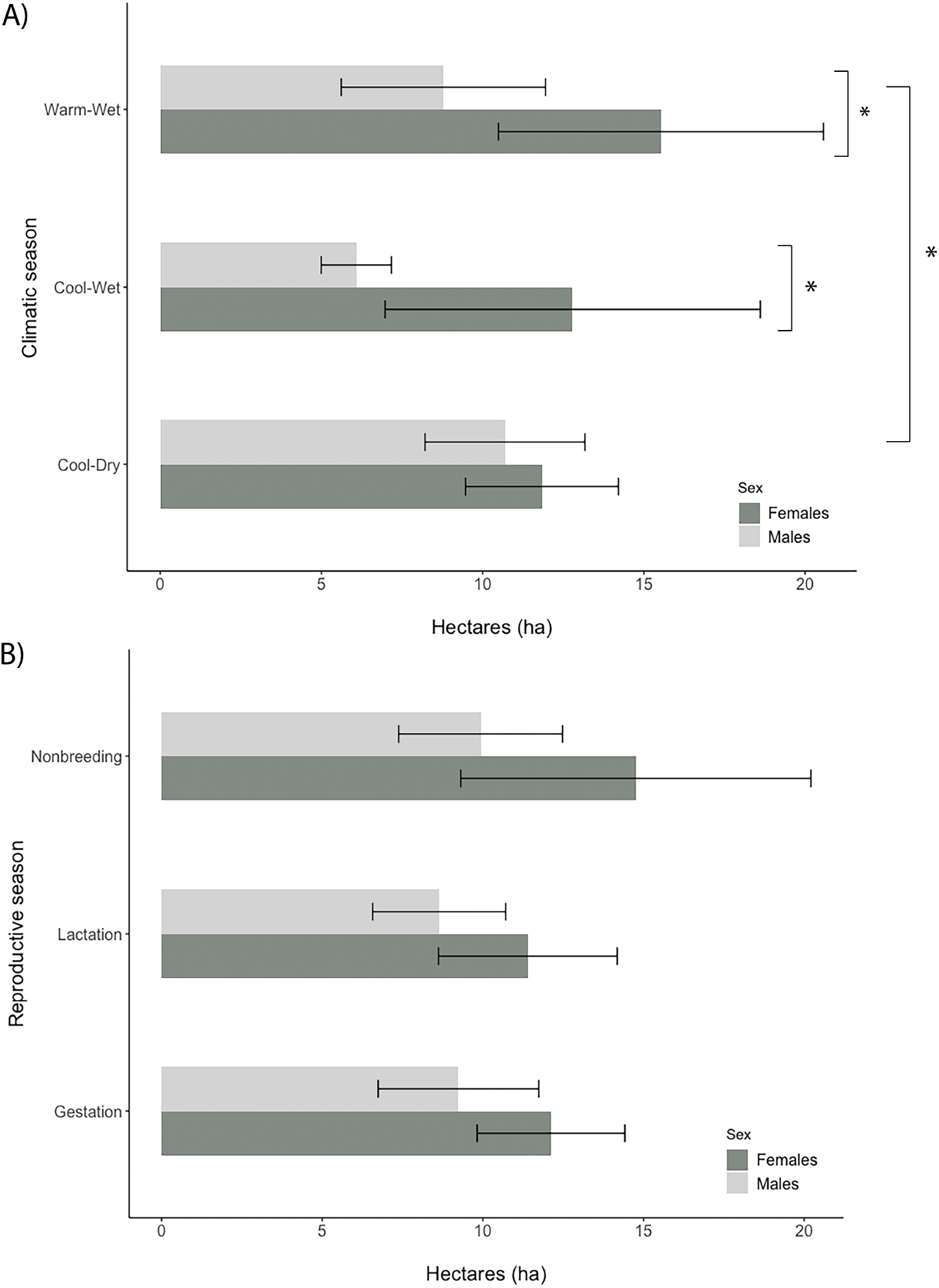
Differences in home range area by sex and both climatic (A) and reproductive (B) seasons. Male range use differed significantly across climatic seasons, using significantly smaller home ranges than females during warm wet and cool wet seasons. Males and females home range size did not differ significantly during the cool-wet period. By contrast, home range size did not differ by sex across reproductive seasons.

**Table 3.**
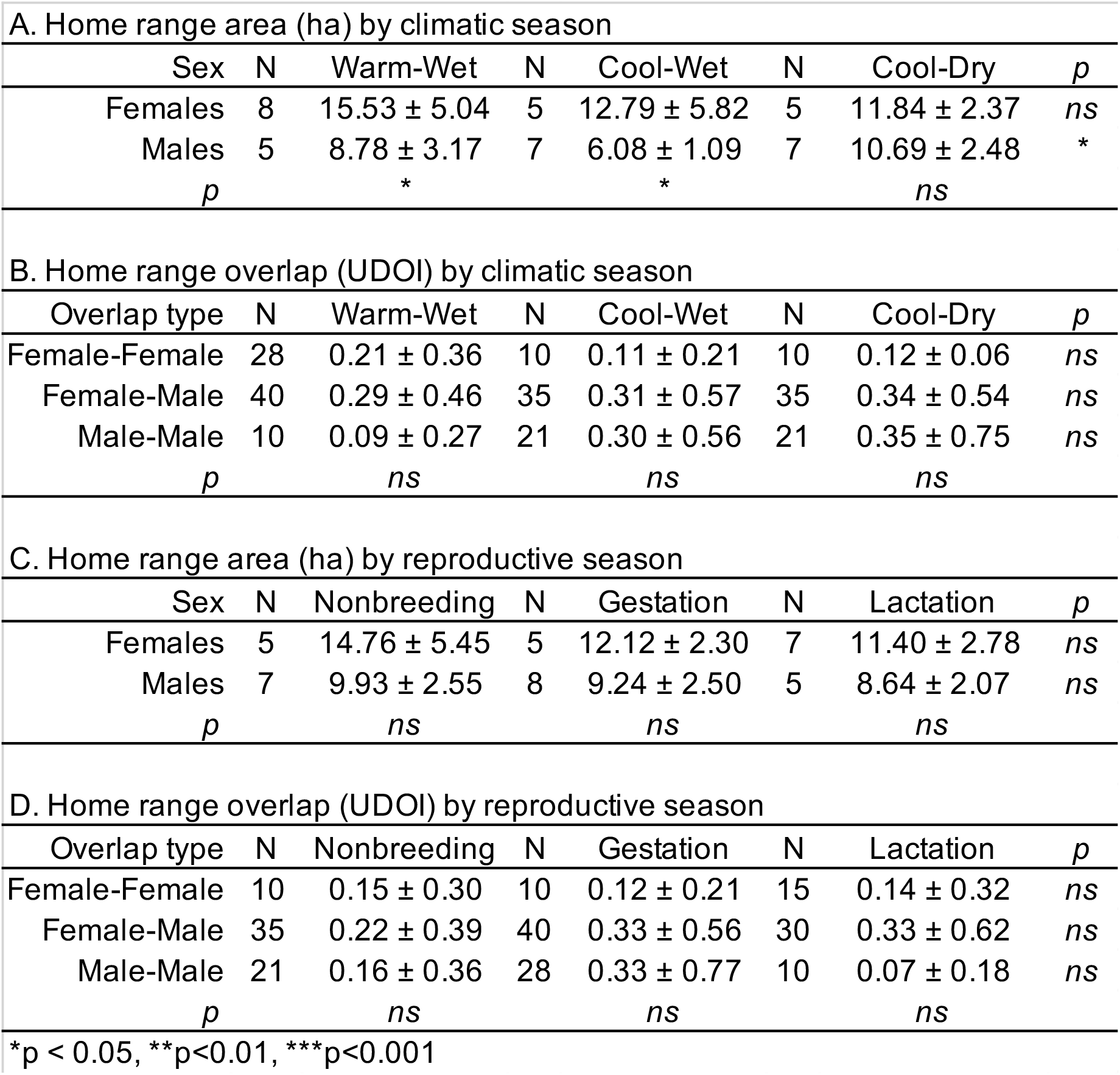
Average seasonal home range area and overlap between sexes. Comparisons include only those individuals with ≥ 10 sampling days per season and home range estimates from at least two of three seasons. Note that statistical comparisons were among only those individuals for which data were available from at least two of the three sampling periods.

| A. Home range area (ha) by climatic season |  |  |  |  |  |  |  |
| --- | --- | --- | --- | --- | --- | --- | --- |
| Sex | N | Warm-Wet | N | Cool-Wet | N | Cool-Dry | *p* |
| Females | 8 | 15.53 ± 5.04 | 5 | 12.79 ± 5.82 | 5 | 11.84 ± 2.37 | *ns* |
| Males | 5 | 8.78 ± 3.17 | 7 | 6.08 ± 1.09 | 7 | 10.69 ± 2.48 | * |
| *p* |  | * |  | * |  | *ns* |  |
| B. Home range overlap (UDOI) by climatic season |  |  |  |  |  |  |  |
| Overlap type | N | Warm-Wet | N | Cool-Wet | N | Cool-Dry | *p* |
| Female-Female | 28 | 0.21 ± 0.36 | 10 | 0.11 ± 0.21 | 10 | 0.12 ± 0.06 | *ns* |
| Female-Male | 40 | 0.29 ± 0.46 | 35 | 0.31 ± 0.57 | 35 | 0.34 ± 0.54 | *ns* |
| Male-Male | 10 | 0.09 ± 0.27 | 21 | 0.30 ± 0.56 | 21 | 0.35 ± 0.75 | *ns* |
| *p* |  | *ns* |  | *ns* |  | *ns* |  |
| C. Home range area (ha) by reproductive season |  |  |  |  |  |  |  |
| Sex | N | Nonbreeding | N | Gestation | N | Lactation | *p* |
| Females | 5 | 14.76 ± 5.45 | 5 | 12.12 ± 2.30 | 7 | 11.40 ± 2.78 | *ns* |
| Males | 7 | 9.93 ± 2.55 | 8 | 9.24 ± 2.50 | 5 | 8.64 ± 2.07 | *ns* |
| *p* |  | *ns* |  | *ns* |  | *ns* |  |
| D. Home range overlap (UDOI) by reproductive season |  |  |  |  |  |  |  |
| Overlap type | N | Nonbreeding | N | Gestation | N | Lactation | *p* |
| Female-Female | 10 | 0.15 ± 0.30 | 10 | 0.12 ± 0.21 | 15 | 0.14 ± 0.32 | *ns* |
| Female-Male | 35 | 0.22 ± 0.39 | 40 | 0.33 ± 0.56 | 30 | 0.33 ± 0.62 | *ns* |
| Male-Male | 21 | 0.16 ± 0.36 | 28 | 0.33 ± 0.77 | 10 | 0.07 ± 0.18 | *ns* |
| *p* |  | *ns* |  | *ns* |  | *ns* |  |

\**p* < 0.05, \*\**p* < 0.01, \*\*\**p* < 0.001

By contrast, there was no discernable variation in spatial community structure for females or males across reproductive seasons (Table 3). Overall, sexes did not differ significantly in home range size (Skillings-Mack test, SM = 1.07, df = 2, *P* = 0.59) or overlap (Skillings-Mack test, SM = 2.04, df = 2, *P* = 0.36) across reproductive seasons, nor did home range size (Skillings-Mack test, SM_males_ = 1.79, df = 2, *P* = 0.41; SM_females_ = 0.400, df = 2, *P* = 0.82) or overlap (Skillings-Mack test, SM_MM_ = 1.66, df = 2, *P* = 0.44; SM_FF_ = 0.95, df = 2, *P* = 0.62; SM_MF_ = 0.50, df = 2, *P* = 0.78) vary significantly within sexes across reproductive seasons.

### Daily range use

Overall, females and males did not differ in their daily path lengths (DPL) (Mann-Whitney *U*=9323.5, *P*=0.926). Females traveled, on average, a daily distance of 1,659.28 m ± 555.21 m whereas males traveled an average of 1,673.83 m ± 598.69 m. However, when included as a fixed effect in our mixed models, we found strong support that DPL was positively related to day length (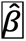 = 0.654, SE=0.111) and varied with reproductive season (Model Weight = 88.1 %; Table 4); the only two models with any support included both of these variables (Table 4). We also found strong support for a sex difference by reproductive season and day length; the model including the sex variable had seven times the support than the model without it (0.881 versus 0.119). We found males to generally move more per day than females (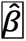 = 11.49, SE=3.79; males = 1, females = 0), with the strongest difference occurring in the lactation season (Table 4). Standardizing to 12 hours of daylight, DPL in both the non-breeding season (female: 1761 ± SE 57.6 m; male: 1948.6 ± SE 122.0 m) and gestation seasons (female: 1736.1 ± 99.48 m; male: 1917.0 ± 296.8 m) were more similar than during lactation, when daily distances traveled by females (840.6 ± SE 106.2 m) were far more restricted than males (1824.6 ± SE 196.7 m).

**Table 4.**
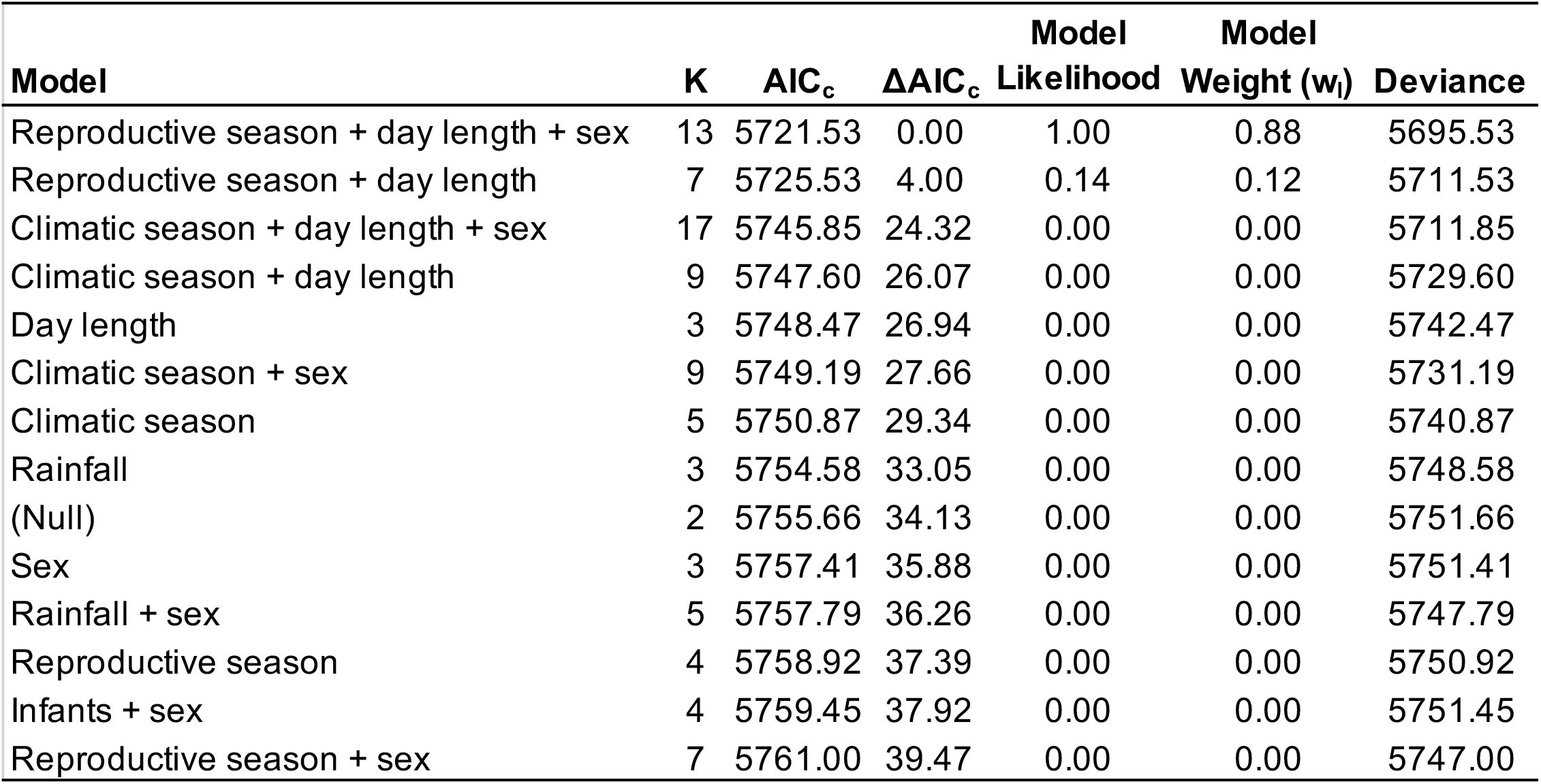
Models using a generalized linear mixed-effects model to estimate daily path length of *Varecia variegata* in the Mangevo community, a primary rainforest site within Ranomafana National Park, Madagascar. Sampling occurred between 6am and 5pm. Fixed effects included day length, daily rainfall (mm), sex, presence of infants, climatic season, and reproductive season. To account for individual variation, ‘individual’ was treated as a random effect. K indicates the number of model parameters.

| Model | K | AIC<sub>c</sub> | ΔAIC<sub>c</sub> | Model Likelihood | Model Weight (w<sub>i</sub>) | Deviance |
| --- | --- | --- | --- | --- | --- | --- |
| Reproductive season + day length + sex | 13 | 5721.53 | 0.00 | 1.00 | 0.88 | 5695.53 |
| Reproductive season + day length | 7 | 5725.53 | 4.00 | 0.14 | 0.12 | 5711.53 |
| Climatic season + day length + sex | 17 | 5745.85 | 24.32 | 0.00 | 0.00 | 5711.85 |
| Climatic season + day length | 9 | 5747.60 | 26.07 | 0.00 | 0.00 | 5729.60 |
| Day length | 3 | 5748.47 | 26.94 | 0.00 | 0.00 | 5742.47 |
| Climatic season + sex | 9 | 5749.19 | 27.66 | 0.00 | 0.00 | 5731.19 |
| Climatic season | 5 | 5750.87 | 29.34 | 0.00 | 0.00 | 5740.87 |
| Rainfall | 3 | 5754.58 | 33.05 | 0.00 | 0.00 | 5748.58 |
| (Null) | 2 | 5755.66 | 34.13 | 0.00 | 0.00 | 5751.66 |
| Sex | 3 | 5757.41 | 35.88 | 0.00 | 0.00 | 5751.41 |
| Rainfall + sex | 5 | 5757.79 | 36.26 | 0.00 | 0.00 | 5747.79 |
| Reproductive season | 4 | 5758.92 | 37.39 | 0.00 | 0.00 | 5750.92 |
| Infants + sex | 4 | 5759.45 | 37.92 | 0.00 | 0.00 | 5751.45 |
| Reproductive season + sex | 7 | 5761.00 | 39.47 | 0.00 | 0.00 | 5747.00 |

## DISCUSSION

Previous work on wild ruffed lemurs has helped to characterize the genus *Varecia* as a taxon notable for its degree of social flexibility; populations exhibit striking variation in group size, social organization and patterns of range use (reviewed in [Vasey, 2003; Baden et al., 2016]. To some extent, this variation can be attributed to sample size, study duration and/or sampling period. However, even in long-term studies (>12 months of behavioral observation) populations have demonstrated remarkable variation in annual and seasonal fluctuations in demographics and behavior (e.g., [Morland, 1991c; Vasey, 2006; Morland, 1991d; Rigamonti, 1993].

Results from this study reveal mixed patterns of ruffed lemur range use, overlap, and daily distances traveled when compared with earlier work (e.g., *Varecia rubra:* [Rigamonti, 1993; Vasey, 2006]; *Varecia variegata:* [Morland, 1991c; Morland, 1991d; Balko, 1998; Ratsimbazafy, 2002]; Table 5). We found that animals travel, on average, nearly 2 km per day. Multiple males and females exploit ranges that together comprise large, communal territories that are more or less spatially distinct from other neighboring social communities. Communal territory size was estimated at approximately 87 hectares using KDE methods, though even with older MCP methods, our estimates fell well within the range of previously reported values (25 to 150 hectares; reviewed in [Vasey, 2003]. Variation between our two size estimates, as well as among ours and earlier studies are largely driven by the methods used to estimate home range area (MCP vs. Kernel Density) and should thus be taken into consideration when comparing range use across studies. Generally, KDEs are more likely to capture the realized spatial distribution compared to MCPs, as they are more robust to outliers, and thus more appropriate for comparison [Powell, 2000; Laver, and Kelly, 2008]. Because ours is the first study to employ the KDE method, we argue that our estimates likely reflect more conservative, and perhaps more biologically meaningful measures of ruffed lemur range use compared to earlier work.

**Table 5.**
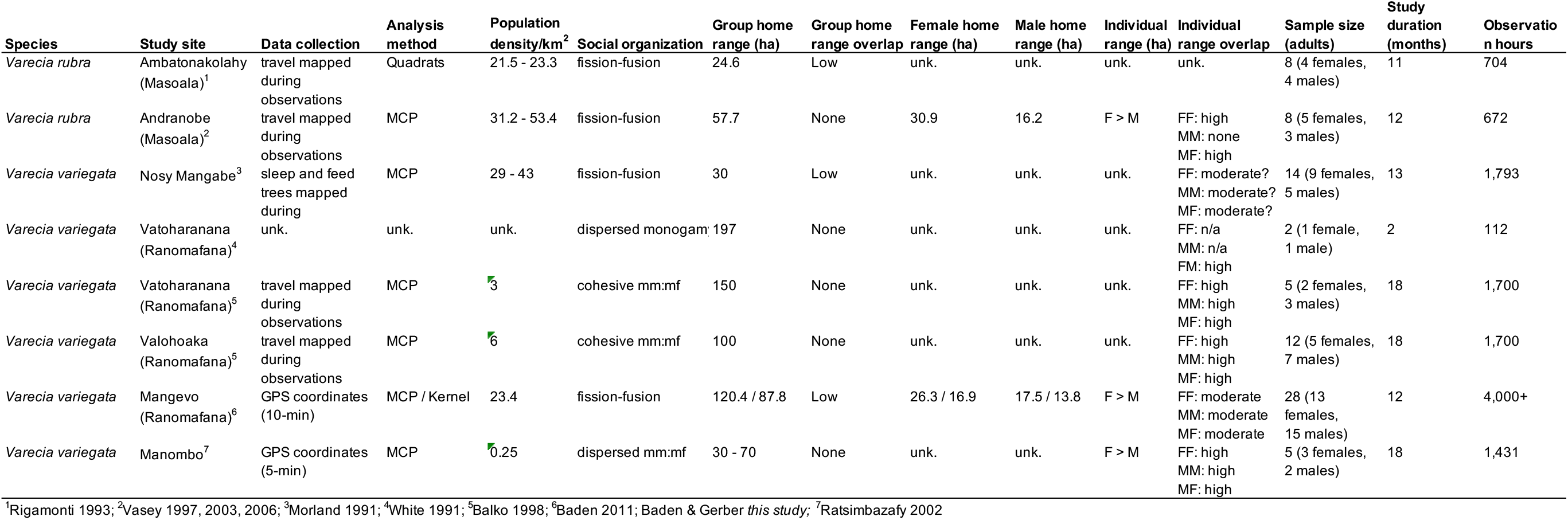
Table comparing results across published ruffed lemur studies.

Neither sex ranged widely throughout the entire communal territory. Rather, both males and females concentrated their movement to only a fraction of the communal range. We report significantly different patterns of range use for males and females; males utilized home ranges that were, on average, 20% smaller than females. These results are largely consistent with sex-segregated patterns of range use described by Morland [1991a,b] and lend support to earlier studies describing females as being central to ruffed lemur social behavior [Morland, 1991a,b; Baden et al., 2016]. Our results differ, however, in that individual ranges were distributed evenly throughout the communal territory and both males and females exhibited moderate levels of annual home range overlap among both same- and mixed-sex dyads. This is in contrast to earlier reports describing males as using small ranges that overlapped extensively with females, but which were discreet from (i.e., non-overlapping with) other neighboring males (e.g., [Vasey, 2006]).

Nevertheless, when considering all available data, broad patterns of ruffed lemur spatial dynamics emerge. It appears that ruffed lemur communities comprise large, multi-male, multi-female groups that live within a communally defended territory, particularly in populations exhibiting high levels of fission-fusion dynamics (e.g., Andranobe, Nosy Mangabe, Kianjavato, Mangevo) [Morland, 1991b; Morland, 1991c; Vasey, 2006; Holmes et al., 2016b; Baden et al., 2016]. Females consistently utilize larger annual home ranges than males, though range use—including DPL— appears to vary both within and between sexes dependent upon climatic season and reproductive state [Morland, 1991c; Morland, 1991d; Vasey, 2006].

### Seasonal variation in range size, overlap and daily distances traveled

Beyond broad characterizations of annual home range size and overlap, we detected significant sex-differences in ranging behavior across seasons. Interestingly, female home range size did not vary significantly by climatic (H1) or reproductive season (H2). Of five females for which we had comparable data across seasons, only one exhibited the expected patterns of home range size variation predicted for seasonal climatic shifts (i.e., largest ranges during warm-wet periods of resource abundance (P1.1); smallest ranges during cool-wet periods of resource scarcity, as per [Vasey, 2006](P1.6); of the remaining four females, two used the relatively largest ranges during the cool-wet (i.e., resource poor) period, while the remaining two females exploited large, and roughly equivalently-sized home ranges in warm-wet (i.e., resource abundant) and cool-dry (i.e., moderate resource) versus cool-wet (i.e., resource poor) seasons (Table 3). Female home range size was equally variable across reproductive seasons (H2) and none met our expectations of home range size (P2.6, 2.8) or overlap (P2.7, 2.10).

Thus, it appears that sex differences in home range size variation were driven primarily by males, whose ranges varied significantly across climatic (H1), but not reproductive seasons (H2). Male range use did not, however, follow expected patterns. We predicted (P1.1) that home range size would be largest during the warm-wet, resource abundant period. Instead, male ranges were largest during the cool-dry period, and directly contradict earlier reports describing seasonal variation in range use as being driven exclusively by females [Vasey, 2006]. Moreover, males consistently traveled longer daily distances than females (~1.9 km vs. ~1.7km/day), and daily female travel was constrained by reproductive state. Female daily path length was consistent across non-breeding and gestation periods (January – September, ~1.7 km/day), but decreased precipitously during the first three months following birth (October – December, ~0.8 km/day). Taken together, our results suggest sex-segregated patterns of range use in ruffed lemurs: females in the Mangevo ruffed lemur community used large, seasonally stable home ranges, but that daily travel was constrained during the lactation period by the presence of dependent offspring. By contrast, males consistently traveled longer daily distances than females, but within smaller, more seasonally flexible areas. While unexpected, we propose a few hypotheses that may help to explain these patterns.

First, in many primate fission-fusion systems, the density and distribution of food resources impact patterns of spatial association (e.g., [Lehmann, and Boesch, 2005]. That female range use in this study did not vary according to climatic seasonality suggests that resources may have been evenly and/or abundantly distributed throughout the communal territory, and that female ranges each may have encompassed high-quality food. Alternatively, given the individual variation we observed, it could be that females in lower quality territories adopted ‘energy minimizing strategies’ by resting more and/or feeding on lower quality food resources until conditions improved (see [Tecot, 2008] for review). Anecdotally, we found asyncrhonous fruit availability across female home ranges, even within the same freeding tree species (Baden, unpublished data), supporting this hypothesis. These and further predictions relating habitat structure and quality to individual patterns of activity and range use in the Mangevo ruffed lemur community are currently underway.

It is also possible that our use of ‘climatic season’ was a poor proxy for resource availability. Although studies—including this one—have found a direct relationship between rainfall and fruit availability in Madagascar [Dewar, and Richard, 2007; Grassi, 2001; Hemingway, 1996; Overdorff, 1993; Meyers, and Wright, 1993], Ranomafana is particularly notable for its highly variable climatic conditions; annual rainfall, phenology and the presence and duration of wet versus dry seasons vary considerably inter-annually [Wright et al., 2005]. Moreover, earlier work in the region (and elsewhere) has demonstrated that relationships between rainfall and fruiting patterns might not always exist [Tecot, 2008; Hemingway, 1998; Van Schaik, and Pfannes, 2005]. Nevertheless, the imperfect relationship between climate and phenology still cannot explain why males—but not females—exhibited seasonal variation in range use. It should be noted that while male range size varied significantly with climatic season, patterns were inconsistent with our expectations. If males were, in fact, altering range use in accordance with resource availability, we might expect larger home range sizes during resource rich months (e.g., [Vasey, 2006]). However, male range size peaked during the cool-dry season, a period of low, but rising resource abundance, suggesting that males may not be modifying spatial patterns according to fruit availability. Instead, we argue that males may, instead, be mapping their ranges onto those of females (i.e., the limiting resource to males), as socioecological theory predicts (e.g., [Sterck et al., 1997]), particularly in the days during the cool-dry season that occur just prior to, during, and following the brief mating period. In support of this hypothesis, we observed higher than average association among males and females according to female reproductive state [Baden et al., 2016]. Thus, it is possible that the observed variation in male home range size is actually a result of males expanding their otherwise small home ranges to map their range use on to those of female associates just prior to the brief mating period in late-June thru early-July.

That male ranging patterns did not vary according to reproductive seasonality is therefore likely an issue of temporal scale in our analysis. Ruffed lemurs, as with other Malagasy primates, are characterized by strict seasonal breeding, and are generally only sexually receptive for two to three days during the year [Brockman et al., 1987; Foerg, 1982; Baden et al., 2013b]. The brevity of estrus in ruffed lemurs therefore complicates seasonal home range estimates according to reproductive state, as it can be difficult to accurately identify the appropriate time frame for analysis. While our behavioral data recorded an influx of males during the “courtship” (pre-receptive, nonbreeding) period (March-June), these changes in behavior were not reflected in our ranging results. In fact, female-male kernel overlap was lowest during the non-breeding season—a period that encompassed this early courtship phase. Unfortunately, we believe that our current temporal scale, which ranged from three to five months per ‘season’, was too large to accurately reflect the shifting patterns of range use across these more fine-scale reproductive stages.

Similarly, we expected patterns of female range use to correspond closely with reproductive state. Ruffed lemurs bear large litters of altricial young, and are notable for their shared nest use and communal infant care, particularly during the first several months after birth [Baden et al., 2013b; Morland, 1993; Pereira et al., 1987; Vasey, 2007]. Because infants cannot cling, and must instead be transported orally by mothers between nests, we expected female movement, and thereby home range size, to be constrained during the period of early lactation and high infant dependence. Moreover, we anticipated female ranges to exhibit greater overlap, a pattern which would have facilitated the communal infant care observed later in infant development (ca. 6-8 weeks; [Baden et al., 2013b]. While patterns of association and daily path length met expectations (i.e., females were significantly less social and daily distances traveled were more constrained during early stages of lactation and high infant dependence [Baden et al., 2016]; this study), home range area and overlap did not differ significantly from other reproductive seasons. Anecdotally, females traveled faster, took more direct routes, and regularly visited diverse areas throughout their range during this time (Baden, unpublished data). Thus, future studies will use newer and more nuanced methods (e.g., ctmm, [Calabrese et al., 2016]; dynamic social network analyses, [Blonder et al., 2012]) to further refine these results.

Finally, ruffed lemurs are ‘boom-or-bust’ breeders, reproducing only during years of resource abundance [Ratsimbazafy, 2002; Baden et al., 2013b]. It may therefore be that patterns of home range area and overlap differ between breeding and non-breeding years, and that the moderate overlap in home ranges observed during this study are actually high relative to years when females do not reproduce. This would also help explain reports of monogamy in the taxon if studies were conducted during non-reproductive years when animals are less social, group members are less cohesive and individuals use smaller, less overlapping home ranges (Baden, unpublished data). Future studies will attempt to document ruffed lemur ranging behaviors across boom and bust years to allow us to test this hypothesis.

### Ruffed lemurs in context

Earlier work has characterized ruffed lemurs as displaying a distinct pattern of fission–fusion dynamics that is both markedly different from and strikingly similar to haplorrhines with fluid fission–fusion grouping patterns [Baden et al. 2016]. Compared to haplorrhines, ruffed lemurs exhibit relatively smaller subgroup size; dramatically lower rates of association; and a female-centered social organization. Adult males and females are equally gregarious, sharing similar numbers of social partners (with the exception of adult male-male dyads [Baden et al. 2016]). Nevertheless, dyadic ruffed lemur social associations are generally sparse and weak, and average relatedness within the community is low [Baden et al. in press]. Our present study further refines this characterization by adding a spatiotemporal component to our understanding of ruffed lemur fission-fusion dynamics. We found that males and females are more-or-less evenly distributed throughout a large, female-defended range, and while female home ranges were larger than males’, group members exhibited equal and moderate home range overlap with other members throughout the community.

Thus, the available data suggest that ruffed lemurs do not adhere to classic ‘male-bonded’ or ‘bisexually-bonded’ community models exhibited by chimpanzees or spider monkeys, two other primates with high fission-fusion dynamics [reviewed in Lehmann and Boesch, 2005; Fig. 1]. (An earlier ‘male-only model was also proposed by Wrangham (1979a), but was later discredited by Pusey (1980).) In ‘male-bonded’ chimpanzee and spider monkey communities, such as those found throughout East Africa (e.g., Gombe: Goodall, 1986; Wrangham et al., 1992; Williams et al., 2002; Kanyawara: Chapman & Wrangham, 1993; Ngogo: Pepper et al., 1999), females are less gregarious than males and utilize relatively small, minimally overlapping or non-overlapping home ranges within a larger, male-defended territory. By contrast, chimpanzee communities in West Africa (e.g., Taï: Lehmann & Boesch, 2005; Lemoine et al. 2019; Bossou, Guinea: Sakura, 1994) and spider monkeys in Yasuni [Spehar et al., 2010] tend to contain males and females that are equally social and range together, with minor ranging differences between sexes due to differential usage of peripheral areas (Lehmann & Boesch, 2005). These differences have been attributed to ecological differences across sites, specifically resource availability and predation pressures.

By contrast, early studies of red-ruffed lemurs from Andranobe described patterns of range use and social association that can best be described as ‘female-bonded’ (Vasey 1997, 2006; Fig. 1). Females were central to social interactions and used large, highly overlapping ranges that encompassed several smaller more-or-less exclusive male ‘core ranges.’ By contrast, in this study, patterns of social and spatial association resembled mixed-sex ‘neighborhoods’. Although females again used larger ranges than males, sexes exhibited equally and evenly overlapping ranges, with neighbors of both sexes affiliating with each other moreso than with other, more distantly located community members (and to the exclusion of members from adjacent communities)[Baden et al. 2016; this study]. In fact, recent work has found that social associations in this species are primarily driven by space use [Baden et al. in press]. Home range overlap significantly predicts the strength of social interactions, moreso than kinship, suggesting that neighbors are more likely to interact than even close relatives. Thus, much like in chimpanzees, ruffed lemurs appear to be flexible in their inter-population patterns of spatial ecology.

Taken together, the unusual sex-segregated patterns of range use and association described herein can be used to characterize ruffed lemurs as exhibiting what we call a ‘neighborhood’ community model (Fig. 1). How these patterns relate to ruffed lemur ecology, reproductive physiology, and evolution remain to be seen. However, several long-term studies are currently working toward addressing these and other questions that will allow us to better understand the complex behavioral ecology of this species.

## ACKNOWLEDGEMENTS

Thanks to ICTE/MICET/Centre ValBio for facilitating research in Madagascar, and ANGAP/Madagascar’s National Parks for permission to conduct research in RNP. Drs. Randy Junge, Felicia Knightly, Angie Simai, Edward E. Louis, Jr. and staff of the Madagascar Biodiversity Project provided invaluable assistance during capture procedures. Reychell Chadwick, Lindsay Dytham, AJ Lowin, Solo Justin, Telo Albert, Lahitsara Pierre, Razafindrakoto Georges, Leroa, and Velomaro provided field assistance, companionship and advice on the ground. Funding was generously provided by National Science Foundation DDIG (ALB: BSC-0725975), J. William Fulbright Foundation (ALB), L.S.B. Leakey Foundation (ALB), Primate Conservation, Inc. (ALB), Primate Action Fund (ALB), Stony Brook University (ALB), and Henry Doorly Zoo (ALB). Research complied with the laws and guidelines set forth by ANGAP/Madagascar National Parks and Stony Brook University’s IACUC (#2005-20081449).

